# Sex- and Subtype-Specific Adaptations in Excitatory Signaling Onto Deep-Layer Prelimbic Cortical Pyramidal Neurons After Chronic Alcohol Exposure

**DOI:** 10.1101/2021.04.30.442127

**Authors:** Benjamin A. Hughes, Todd K. O’Buckley, Giorgia Boero, Melissa A. Herman, A. Leslie Morrow

**Author notes:** **CORRESPONDING AUTHOR:** Benjamin A. Hughes, Ph.D., Mailing Address: University of North Carolina at Chapel Hill, 3027 Thurston Bowles, CB 7178, Chapel Hill, NC 27599.

## Abstract

Long-term alcohol use results in behavioral deficits including impaired working memory, elevated anxiety, and blunted inhibitory control that are associated with prefrontal cortical (PFC) dysfunction. Preclinical observations demonstrate multiple impairments in GABAergic neurotransmission onto deep-layer principal cells (PCs) in prelimbic cortex that suggest dependence-related cortical dysfunction is the product of elevated excitability in these cells. Despite accumulating evidence showing alcohol-induced changes in interneuron signaling onto PCs differ between sexes, there is limited data explicitly evaluating sex-specific ethanol effects on excitatory signaling onto deep-layer PCs that may further contribute to deficits in PFC-dependent behaviors. To address this, we conducted electrophysiological and behavioral tests in both male and female Sprague-Dawley rats to evaluate the effects of chronic ethanol exposure. Among our observations, we report a marked enhancement in glutamatergic signaling onto deep-layer PCs in male, but not female, rats after alcohol exposure. This phenomenon was furthermore specific to a sub-class of PC, sub-cortically projecting Type-A cells, and coincided with enhanced anxiety-like behavior, but no observable deficit in working memory. In contrast, female rats displayed an alcohol-induced facilitation in working memory performance with no change in expression of anxiety-like behavior. Together, these results suggest fundamental differences in alcohol effects on cell activity, cortical sub-circuits, and PFC-dependent behaviors across male and female rats.

## INTRODUCTION

Alcohol use disorders (AUDs) are a major public health hazard, with AUDs contributing to ∼6% of preventable fatalities worldwide and costing nearly 3% of the gross domestic product in developed countries [1, 2]. Though of great concern, AUDs encompass a wide range of aberrant drinking behaviors that elicit myriad pathological peripheral and central nervous system adaptations, and the discrete mechanisms by which these behaviors manifest are therefore difficult to discern. Of particular interest to our laboratory is the pathogenesis of dependence-related alcohol consumption, as dependent individuals, though comprising only ∼5-14% of alcohol abusers, account for the majority of the public health burden stemming from AUDs [3, 4]. Excessive alcohol consumption characteristic of dependence is specifically associated with diminished executive control of intake as well as elevated anxiety experienced during acute withdrawal that drives consumption [5]. These observations consistently correlate with notable structural and functional impairments in prefrontal cortical (PFC) function, leading models of addiction to include PFC dysfunction as a core component [6].

Nevertheless, much of the literature investigating alcohol effects on the PFC neglect to include females, despite comparable lifetime risks for developing AUDs among men and women (∼36% and 22.7%, respectively [7]) as well as recent policy directives by the National Institutes of Health to include female subjects [8]. Indeed, the assumption that AUDs manifest comparably between sexes conflicts with available clinical and preclinical evidence demonstrating key differences in both the acquisition and progression of AUDs, finding that males escalate alcohol consumption sooner than females [9-11].

Our laboratory has used rat models to explore the molecular adaptations to chronic ethanol exposure that are associated with dependence-like symptoms such as increased seizure susceptibility and anxiety-like behavior, finding latent sex differences in responses to alcohol. For instance, we have previously reported dimorphic alterations in GABA_A_ receptor subunit expression, reporting that, while α1 subunit expression is reduced in both males and females following chronic alcohol exposure [12, 13], α3 subunit expression is altered only in males [14]. Furthermore, elevations in N-methyl-D-aspartate (NMDA) receptor function and expression that promote increased cell excitability are reliably observed with chronic alcohol exposure in male rats [15, 16] also appear to be sex-specific [17].

More recently, we and others have specifically investigated alcohol effects on the functional properties of cortical neurons between sexes, finding important similarities and differences [18-20]. Specifically, in seeking to localize ethanol-induced GABA_A_ receptor subunit expression changes in cortex, our laboratory reported equivalent reductions in GABAergic spontaneous inhibitory post-synaptic currents (sIPSCs) onto deep-layer principal cells (PCs), the primary source of efferent signaling in PFC, after alcohol exposure in male and female rats. Reductions in sIPSC frequency furthermore occurred in tandem with post-synaptic reductions in α1 subunit expression, consistent with elevated PC excitability during acute withdrawal [12, 13]. Interestingly, though, interneuron recordings revealed that while fast-spiking interneurons displayed equivalent reductions in excitability in both sexes consistent with reductions in sIPSC frequency, Martinotti interneurons exhibited markedly divergent responses in excitability between males and females [19]. It is unclear how such differential changes in intra-cortical feedback inhibition affect local microcircuit function, and raise an important question as to whether other functional adaptations induced by alcohol exposure in PFC are similarly sex-dependent. Building on these observations of GABAergic signaling on deep-layer PCs we therefore hypothesized that glutamatergic tone onto these cells from superficial layer PCs would display similarly sex-specific responses to chronic ethanol exposure that further promote pyramidal cell hyperexcitability. In addition, we sought to probe whether potential functional changes correlated with alterations in PFC-dependent behavior, finding that chronic alcohol exposure elicits complex functional and behavioral adaptations that do not occur uniformly between sexes.

## MATERIALS AND METHODS

### Animal Care and Exposure Paradigm

Male and female Sprague Dawley rats were obtained from in-house breeding using breeders sourced from Envigo (Indianapolis, IN) and pair-housed in a temperature and humidity controlled vivarium under a 12 hour light/dark cycle and given free access to water and food unless otherwise noted. All animals were PN90-130 at time of testing, and all procedures were carried out in accordance with guidelines specified by the Institutional Animal Care and Use Committee at the University of North Carolina at Chapel Hill. Ethanol exposure consisted of a once-daily dose of ethanol (5.0 g/kg, 25% v/v) or water via intragastric gavage (i.g.) approximately 2 hours into the light cycle that reliably elicits BECs in both male and female rats of ∼250-300 mg/dl ([13]; **Supplemental Figure S1**). Experimenters performing gavage possessed extensive training with the technique and when performed correctly is not especially stressful, eliciting comparable EPM observations as other forced-administration models [21, 22]. Animals received 15 consecutive days of exposure, all experiments performed 24 hours after the last exposure, and behavioral and physiological experiments performed in separate animals. At the conclusion of all experiments, rats were sacrificed by rapid decapitation and brains collected for electrophysiological or biochemical analyses.

### Elevated Plus Maze (EPM) and Delayed Non-Match-to-Sample (NMS) T-Maze

EPM was conducted 24 hours after the last water/ethanol exposure in a partially lit room (∼20 lux), using previously described methods [12]. The Delayed NMS T-maze task was performed according to previously described procedures [23, 24]. Detailed description of behavioral tasks and procedures can be found in **Supplementary Materials**.

### Electrophysiological Recordings

After sacrifice, brains were extracted and ∼300 µM thick coronal slices containing the mPFC were prepared in oxygenated ice-cold sucrose solution (in mM: 200 sucrose, 1.9 KCl, 1.2 NaH2PO4, 6 MgCl2, 0.5 CaCl2, 0.4 ascorbate, 10 glucose, 25 NaHCO3, osmolarity adjusted to ∼310 mOsm) using a Leica VT1000S vibratome (Buffalo Grove, IL). Slices were then transferred to a continuously oxygenated holding chamber containing standard aCSF solution (in mM: 125 NaCl, 2.5 KCl, 1.5 NaH2PO4, 1.3 MgCl2, 2.6 CaCl2, 0.4 ascorbate, 10 glucose, 25 NaHCO3, osmolarity adjusted to ∼310 mOsm) warmed to 32-34° C and incubated for 30min, after which the holding chamber was allowed to cool to room temperature for 1 hour. Whole-cell recordings of prelimbic Layer V pyramidal cells were performed at room temperature using an Axon Instruments Multiclamp 700A amplifier sampling at 10 kHz and filtered at 2 kHz. Recordings were performed in standard aCSF solution supplemented with 50 µM dl-AP5 and 10 µM CGP-52432, and using 2-4 mOhm borosilicate glass filled with a K-gluconate-based internal solution (in mM: 120 K-gluconate, 10 KCl, 2 MgCl2, 5 EGTA, 10 HEPES, 3 Na-ATP, 0.5 Na-GTP, 2 phosphocreatine, pH 7.4, osmolarity adjusted to ∼290 mOsm with sucrose). Following gigaseal formation and breakthrough, cells were held at -70 mV and electrically-evoked excitatory post-synaptic currents (eEPSCs) were obtained via a bipolar stimulating electrode placed in Layers I-III of prelimbic cortex. Stimulation intensity was tuned to elicit a ∼200-300 pA response, and baseline stimulation frequency-response recordings obtained. Additional description of electrophysiological methods can be found in **Supplementary Materials**.

### Statistical Analysis

Data are expressed as mean±SEM unless otherwise noted. Electrophysiological data were analyzed using linear mixed modeling and Satterthwaite approximation of effective degrees of freedom to model extraneous inter-animal variability as a random-effect with SAS 9.4 software (SAS Institute; Cary, NC) as previously reported [19, 25]. Behavioral data were analyzed by unpaired t-tests with Welch’s correction (EPM) and two-way ANOVA with Bonferroni post-hoc tests (NMS T-maze) using GraphPad Prism 6.0 (San Diego, CA) and statistical power estimated using the free software GPower 3.1. As direct statistical comparison of sex would require prohibitively high *n*’s for the present experiments, male and female datasets were analyzed independently. For all analyses significance is defined as p<0.05 (*<0.05, **<001, ***<0.001).

## RESULTS

### Sex-Specific Enhancement of Excitatory Signaling onto Deep Layer Pyramidal Neurons After Ethanol Exposure

While post-synaptic changes in glutamatergic and GABAergic signaling onto deep-layer PCs during acute withdrawal have been described [13, 16, 26], it is unclear whether chronic alcohol exposure affects Layer II/III excitatory signaling onto deep-layer PCs, or whether such effects occur in both sexes. Therefore, we evaluated post-synaptic responses in deep-layer PCs to repeated electrical stimulations of superficial layer PCs, as this is a well-validated means of gauging presynaptic glutamate release probability [27-29]. As shown in **Figure 1**, alcohol exposure shifted post-synaptic responses to electrical stimulation from a facilitating profile towards repeated-pulse depression at both low-(3 Hz; **Figure 1A**) and high-frequencies (30 Hz; **Figure 1B**) in male rats, indicative of elevated glutamate release probability [3 Hz: F_(3,132)_Interaction=8.87, p<0.0001; F_(3,132)_Stimulation=8.16, p=<0.0001; F_(1,19.4)_Treatment=22.55, p=0.0001: 30 Hz: F_(1,132)_Interaction=3.05, p=0.031; F_(3,132)_Stimulation=16.62, p<0.0001; F_(1,19.5)_Treatment=4.8, p=0.041]. Female rats, however, displayed no apparent effect of chronic ethanol on repeated-pulse responses (RPR) at either frequency [**Figure 1D-E**: 3 Hz: F_(1,14.5)_Treatment=2.39, p=0.143: 30 Hz: F_(1,16.2)_Treatment=0.06, p=0.807]. The observation that this effect occurs irrespective of initial stimulation intensity further suggests a generalized enhancement of presynaptic release probability (**Figure 1C,F**). Additionally, application of picrotoxin (100 µM) did not alter responses, showing that changes in presynaptic release are likely not the product of local inhibitory processes [**Figure 1G-I**: F_(1,8.07)_Treatment=10.3, p=0.012; F_(1,151)_Picrotoxin=1.91, p=0.17].

**Figure 1:**
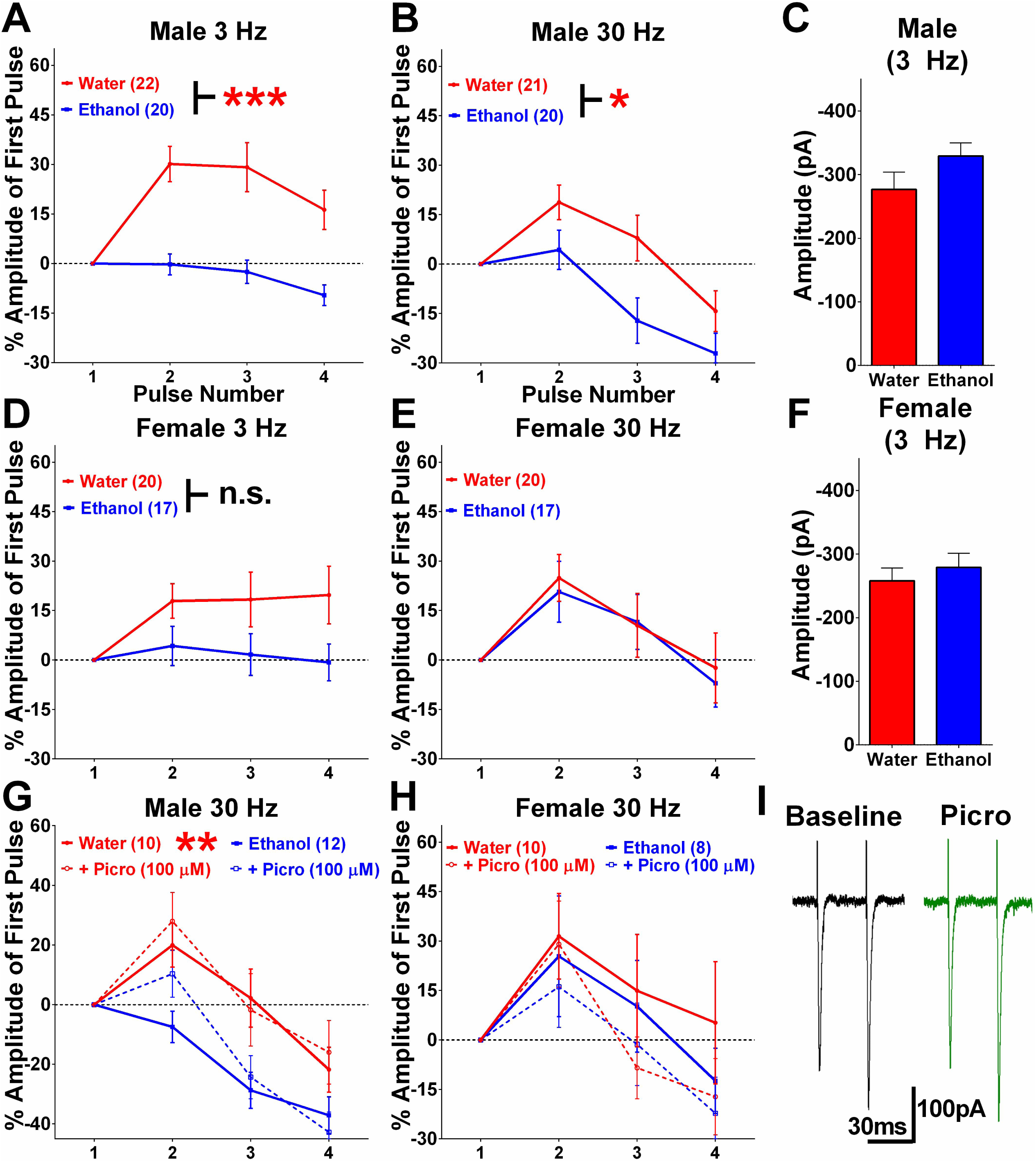
Chronic alcohol exposure elevates superficial layer excitatory signaling onto deep-layer pyramidal neurons in prelimbic PFC. **A-B**) Chronic alcohol exposure shifts electrically-evoked AMPA receptor-mediated currents responses from repeated-pulse facilitation towards depression in male rats across a range of stimulation frequencies [3 Hz: F_(1,19.4)_ Treatment=22.55; 30 Hz: F_(1,19.5)_ Treatment=4.8; n=20-22 cells from 10-12 rats/group]. **C**) Differential repeated-pulse responses (RPR) between groups are not the product of altered sensitivity to electrical stimulation. **D-E**) No effect of chronic ethanol exposure is observed on RPR in female PCs [3 Hz: F_(1,14.5)_ Treatment=2.39; 30 Hz: F_(1,16.2)_ Treatment=0.06; n=17-20 cells from 9-11 rats/group]. **F**) Similar to males, there was no apparent change in sensitivity of electrical stimulation between exposure groups. **G-H**) Application of picrotoxin (100 µM) did not alter RPR in either sex, suggesting this phenomenon is not activity-dependent. **I**) Representative traces depicting electrically-evoked (30 Hz) AMPA currents pre- and post-application of picrotoxin.

### Heterogeneity in Alcohol Effects on Principal Cell Excitability

Despite a relatively robust *n*, we observed substantial variability in RPRs in both males and females that suggested the possibility of a non-homogeneous cell population, and indeed distinct subtypes of deep-layer PCs have previously been described [30-33]. These cell subtypes, termed Type-A and Type-B, are readily discriminable by physiological criteria including the presence of a robust hyperpolarization-induced voltage sag (V_H_; **Figure 2A**, red arrows; **Figure 2B-C**), differential facilitating responses to presynaptic stimulation (**Figure 2D-E**; representative traces shown in **Figure 2F**), and intrinsic membrane characteristics (**Supplemental Table 1**). Additional experiments were therefore conducted, revealing that subtyping according to these criteria (e.g. total V_H_ >11.0 mV, facilitating RPR profile, and C_Mem_ >95.0 pF) reliably distinguished Type-A from Type-B cells. As shown in **Figure 2G**, both cell types were observed in approximately equal frequency in male and female rats, and irrespective of exposure group. Thus, we next asked whether alcohol-induced changes in presynaptic glutamate release from superficial-layer PCs may be preferential for either PC subtype.

**Figure 2:**
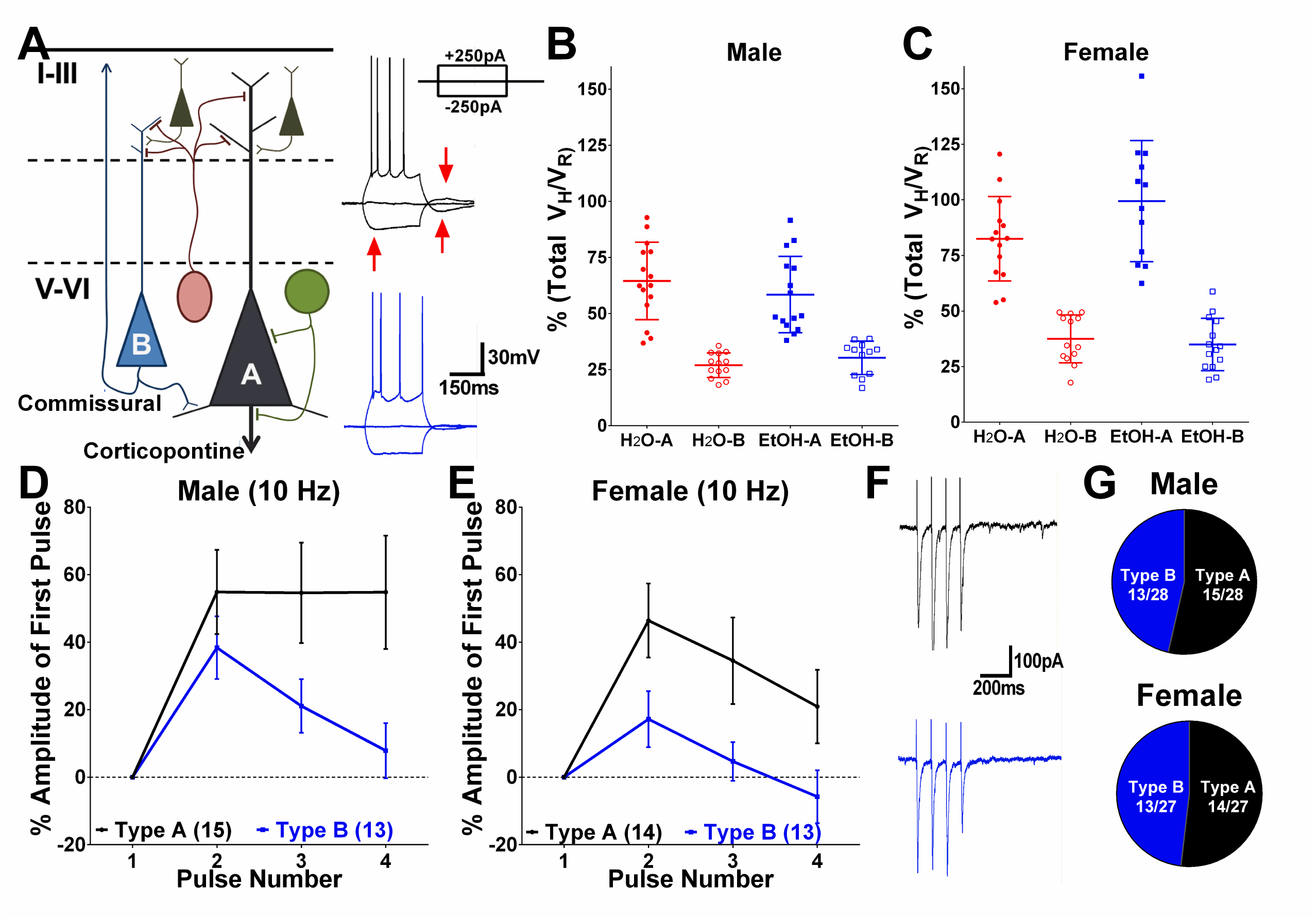
Deep-layer PCs exhibit distinct subtypes in both male and female rats. **A)** Simplified circuit diagram showing differential innervation patters of Type-A and -B PC subtypes, as well as representative current-clamp traces from each. Red arrows show presence of HCN-mediated voltage characteristic of Type-A cells. **B-C)** Mean V_H_ (± S.D.) between cell subtypes in males and females [males: Type-A Water=64.5±17.3; Type-A EtOH=58.4±17.1; Type-B Water=27.0±5.5; Type-B EtOH=30.3±7.4: females: Type-A Water= 82.5±19.0; Type-A EtOH=99.5±27.2; Type-B Water=37.5±10.7; Type-B EtOH=35.0±11.8]. Females displayed somewhat greater V_H_ in all cells compared to males. **D-E)** Type-A and –B cells exhibit differentiable repeated-pulse responses to electrical stimulation in both male and female rats. **F)** Representative voltage-clamp traces showing repeated-pulse responses between cell subtypes. **G)** PC subtypes were observed in approximately equal frequency in male and female rats.

As shown in **Figure 3A**, we observe a significant shift from facilitation towards RPR depression after alcohol exposure in Type-A PCs of male rats that is not observed in females (**Figure 3B**) [male: F_(3,100)_Stimulation=10.2, p<0.0001; F_(1,8.84)_Treatment=5.58, p=0.0344: female: F_(3,80.1)_Stimulation=6.28, p=0.0007; F_(1,7.08)_Treatment=0.04, p=0.853]. Based on previous reports that alcohol-induced changes in presynaptic release are calcium dependent [11, 34], we hypothesized that reducing extracellular calcium would reduce release probability, observed as a shift towards RPR facilitation, to a greater degree in alcohol-exposed PCs. **Figures 3C-E** demonstrate that reducing extracellular calcium elicits a significant enhancement in RPR facilitation in both male and female Type-A cells [males: F_(1,127)_CalciumMod=93.4, p<0.0001: females: F_(1,175)_CalciumMod=81.26, p<0.0001] to equivalent levels between water/ethanol-exposed groups. Analyzing the % change in RPR between conditions, however, revealed significantly higher modulation in male ethanol-exposed Type-A cells (**Figure 3D**), suggesting alcohol effects on presynaptic release in males involve a calcium-dependent mechanism [males: F_(1,13.2)_Treatment=4.92, p=0.0473]. We further measured current-evoked spiking, and alcohol exposure resulted in a significant left-ward shift (e.g. >100% increase at 150 pA) in intrinsic activity in male, but not female, Type-A cells that further underscores the ethanol-induced bias toward hyperexcitability in this population [**Figure 3G-H**; males: F_(1,13.2)_Treatment=4.92, p=0.0446: females: F_(1,10.4)_Treatment=0.11, p=0.744].

**Figure 3:**
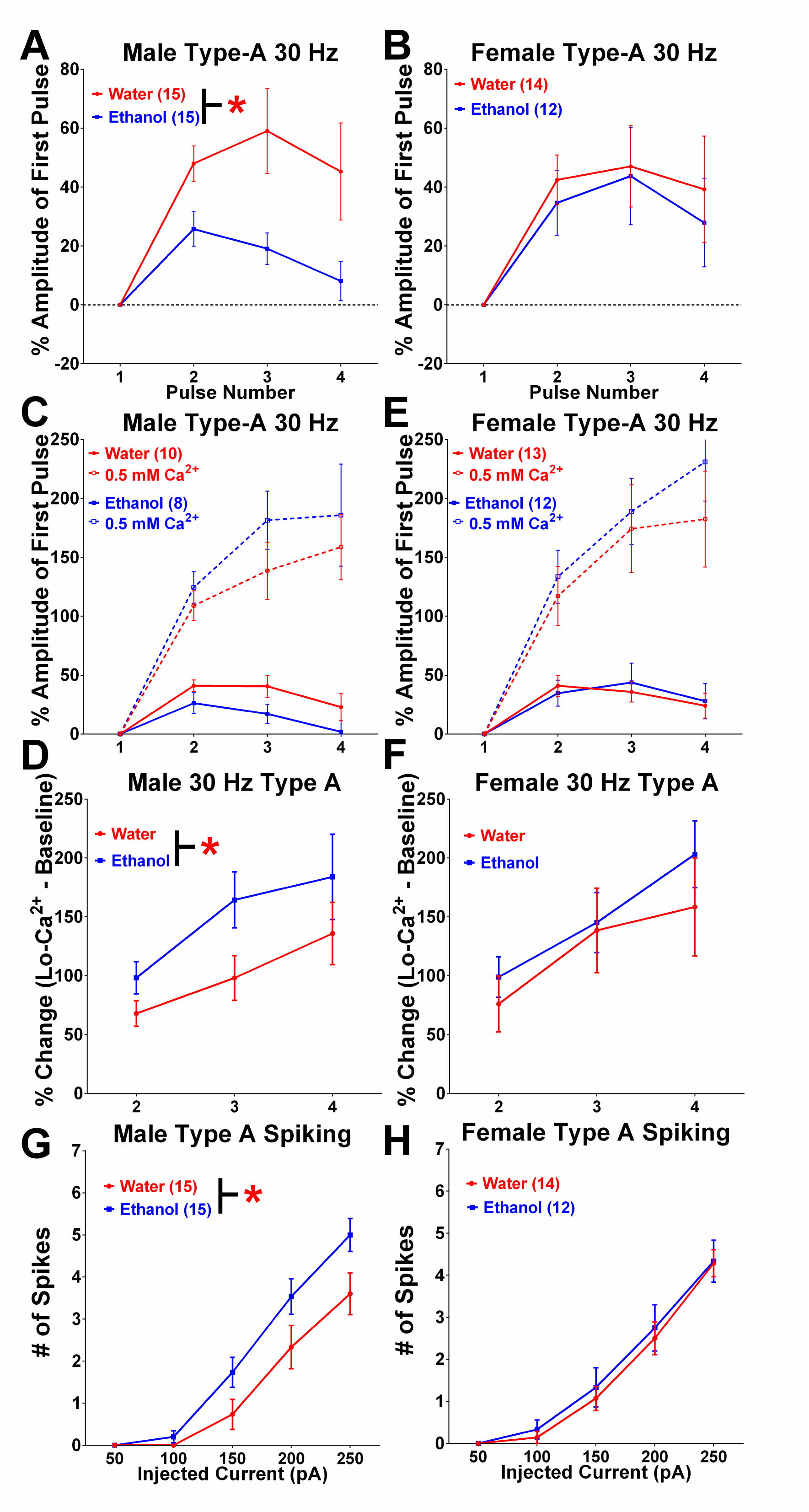
Sex-specific alcohol enhancement of glutamate release probability onto Type-A PCs. **A-B**) Type-A cells in males display alcohol-induced shift toward repeated-pulse depression that is not observed in female rats, consistent with observations in **Figure 1** [males: F_(1,8.84)_ Treatment=5.58, p=0.0344; n=15 cells from 7-8 rats/group: females: F_(1,7.08)_ Treatment=0.04, p=0.853; n=12-14 cells from 6-7 rats/group]. **C-D**) Reducing extracellular calcium concentration reduces glutamate release probability, shown as a profound elevation in repeated-pulse facilitation [F_(1,127)_ CalciumMod=93.4, p<0.0001; n=8-10 cells from 5-6 rats/group]. Significantly larger pulse facilitation after modulating calcium concentration in ethanol-exposed cells is indicative of alcohol-induced enhancement of glutamate release probability on these neurons [F_(1,13.2)_ Treatment=4.92, p=0.0473]. **E-F**) Females display similarly enhanced pulse facilitation in low-calcium solution, with no difference observed between water- and alcohol-exposed groups [F_(1,175)_ CalciumMod=81.26, p<0.0001; n=12-13 cells from 6-7 rats/group]. **G-H**) Alcohol exposure results in significantly elevated excitability of Type-A cells in male, but not female, rats [males: F_(1,13.2)_ Treatment=4.92, p=0.0446; n=15 cells from 7-8 rats/group].

Evaluation of RPRs in Type-B cells revealed no apparent differences between water and ethanol-treated groups in either male or female rats (**Figure 4A-B**) [males:F_(1,9.5)_ Treatment=0.25, p=0.63; F_(2,38.9)_Stimulation=4.42, p=0.001: females: F_(1,9.2)_Treatment=0.99, p=0.35; F_(3,87.3)_Stimulation=6.36, p=0.0006]. Reducing extracellular calcium similarly elevated RPR in both male and female rats to equivalent levels (**Figure 4C,E**) [males: F_(1,131)_CalciumMod=49.71, p<0.0001; F_(1,10.1)_Treatment=0.40, p=0.54: females: F_(1,112)_CalciumMod=55.72, p<0.0001; F_(1,5.3)_Treatment=0.29, p=0.61], with no observable difference in % change between conditions (**Figure 4D,F**) [males: F_(1,9.5)_Treatment=0.48, p=0.506: females: F_(1,4.9)_Treatment=1.67, p=0.25]. Consistent with minimal ethanol effects on these cells, we further observed no apparent difference between exposure groups on intrinsic excitability in either sex (**Figure 4G-H**) [males: F_(1,9.8)_Treatment=0.37, p=0.555: females: F_(1,12.3)_Treatment=0.72, p=0.411].

**Figure 4:**
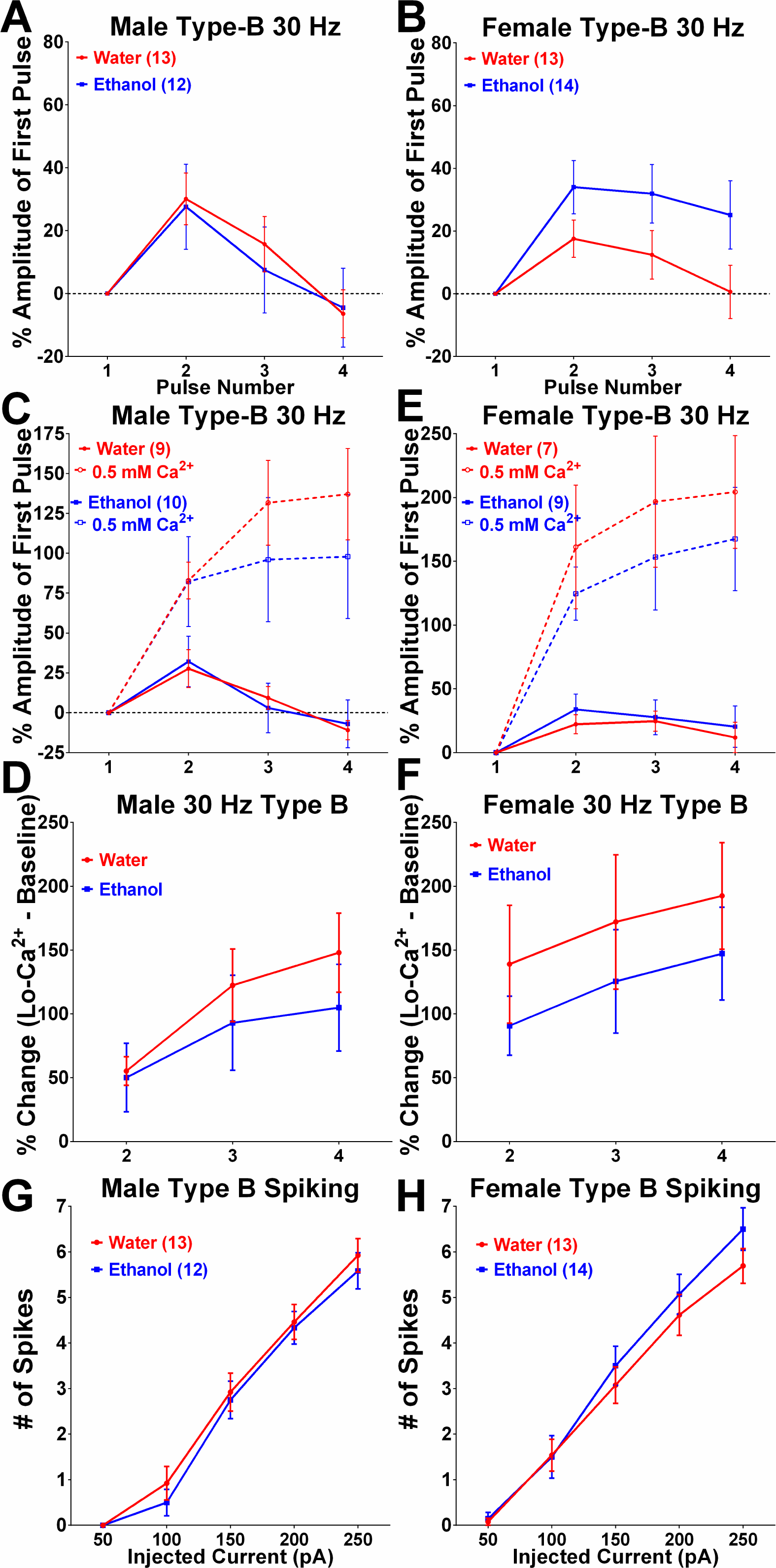
Alcohol exposure does not alter glutamate release onto Type-B PCs in male or female rats. **A-B)** Type-B PCs do not exhibit any appreciable change in repeated-pulse responses after chronic ethanol exposure in either male or female rats (males; n=12-13 cells from 6-7 rats/group: females; n=13-14 cells from 6-7 rats/group). **C-F)** Modulating extracellular calcium concentration results in an expected shift toward pulse facilitation in both sexes, with no difference between water- and alcohol-exposed groups [males: F_(1,131)_ CalciumMod=49.71, p<0.0001; n=9-10 cells from 6-7 rats/group: females: F_(1,112)_ CalciumMod=55.72, p<0.0001; n=7-9 cells from 4 rats/group]. **G-H)** Alcohol exposure does not alter excitability of Type-B cells in either male or female rats.

### Differential Effects of Alcohol Exposure on PFC-related Behaviors by Sex

The vast interconnectivity of the PFC permits this region to influence a wide variety of behaviors. As such, altering prelimbic PFC signaling elicits a number of cognitive impairments including deficits in working memory [23, 35] and the expression of anxiety-like behaviors [11, 12, 34, 36]. On this basis, we therefore sought to determine whether the observed subtype-specific changes in deep-layer PCs after chronic alcohol exposure were associated with specific changes in behavior. To gauge working memory, we employed the Delayed NMS T-maze task, training rats for 7 consecutive days followed by 15 days of water/ethanol exposure and then 3 days of testing (**Figure 5A**). Both male and female rats reached a satisfactory level of performance on the task by the end of training, and no significant differences in training between groups were observed (**Figure 5-B,E**). While increasing delay time between forced-arm and choice trials resulted in an expected decrease in choice accuracy, male rats displayed no apparent difference in performance between water- and alcohol-exposed groups, either at 24 hr acute withdrawal (**Figure 5C**) or over 3 days (**Figure 5D**) of cumulative testing [24 hr test two-way ANOVA: F_(2,56)_Time=7.38, p=0.0014; F_(1,28)_Treatment=0.0438, p=0.836: 3 day test two-way ANOVA: F_(2,56)_Time=22.6, p<0.0001; F_(1,28)_Treatment=0.732, p=0.399]. Female rats, however, displayed a significant interaction between alcohol exposure and delay time at 24 hr acute withdrawal (**Figure 5F**) [two-way ANOVA, Bonferroni post-test: F_(2,36)_Interaction=5.609, p=0.0076; F_(2,36)_Time=10.3, p=0.0003; F_(1,18)_Treatment =4.20, p=0.0553] and a robust main effect of alcohol exposure over 3 days (**Figure 5G**) of cumulative testing [two-way ANOVA, Bonferroni post-test: F_(2,36)_Interaction=4.249, p=0.022; F_(2,36)_Time=35.97, p<0.0001; F_(1,18)_Treatment=10.84, p=0.004] that indicate ethanol exposure improved performance of female rats on this task.

**Figure 5:**
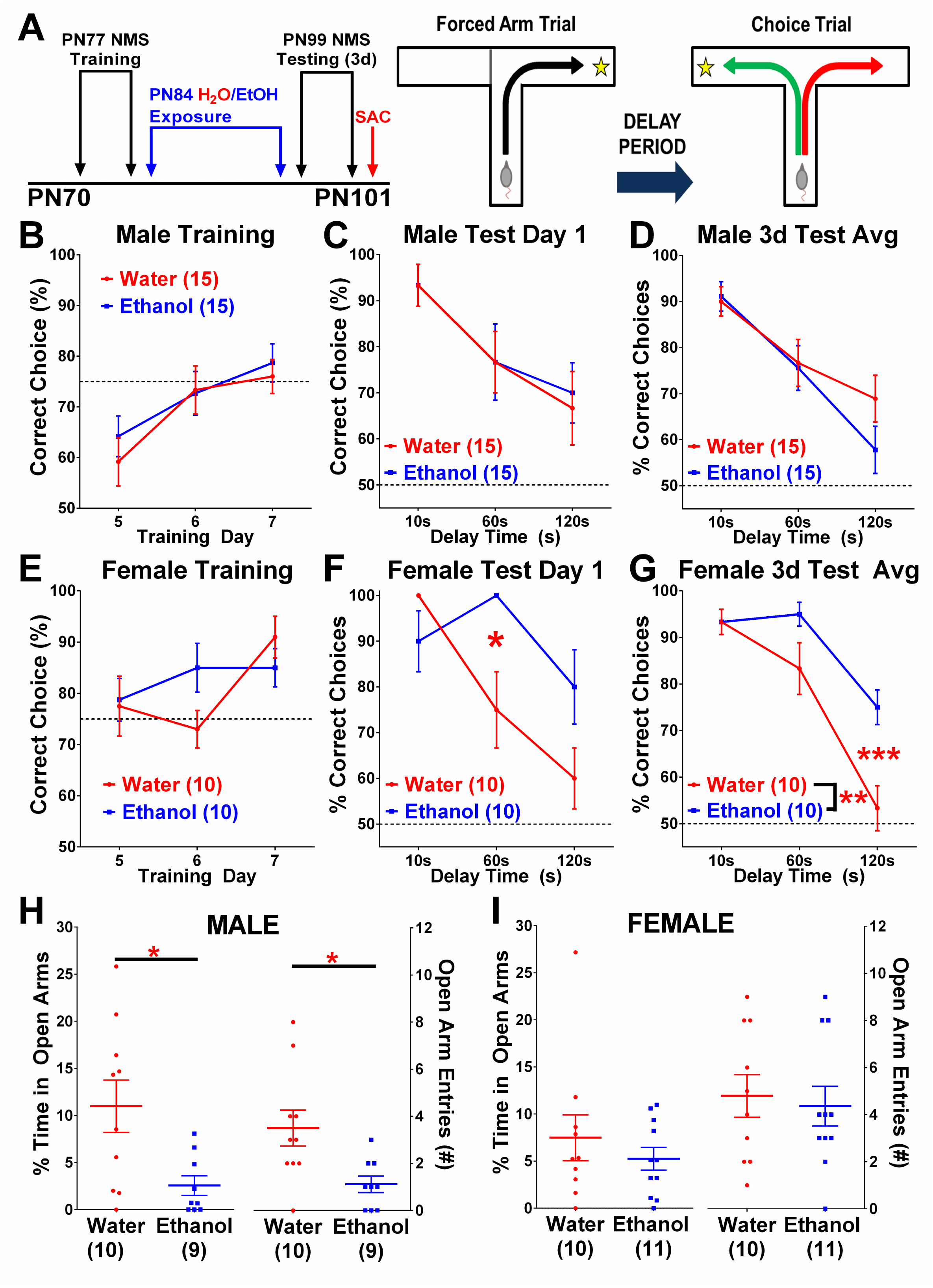
Chronic ethanol exposure alters PFC-dependent behaviors in a sex-dependent manner. **A**) Training and exposure paradigm for the delayed non-match-to-sample (NMS) T-maze task. **B**) Male water- and ethanol-exposed rats perform equally during task training prior to alcohol exposure. **C-D**) Increasing delay between forced and choice trails results in expected reductions in task performance that does not differ between exposure groups at 24 hr acute withdrawal and is consistent across 3 consecutive test days. **E**) Similar to males, female groups perform equivalently during task training prior to exposure. **F-G**) Alcohol-exposed female rats exhibit significantly higher performance on the delayed NMS T-maze task at 60 seconds delay compared to water-exposed rats during 24 hr acute withdrawal [F_(2,36)_ Interaction=5.609, p=0.0076; Bonferroni post-test]. Cumulative performance across 3 consecutive test days reveals a robust enhancement in task performance after alcohol exposure [F_(1,18)_ Treatment=10.84, p=0.004]. **H**) Chronic alcohol exposure in male rats results in significant elevations in anxiety-like behavior on an elevated plus maze [Open Arm Time t_11.5_=2.841, p=0.0155; Open Arm Entries t_12.6_=2.842, p=0.0143]. **I**) In contrast, female rats display no apparent effect of ethanol exposure on expression of anxiety-like behavior.

In contrast, measurement of anxiety-like behavior revealed significant effects in males, but not females. Specifically, male ethanol-exposed rats displayed significant reductions in both the percent time in open-arm and open-arm entries on EPM (**Figure 5H**), consistent with elevated anxiety during 24 hr acute withdrawal [unpaired two-tailed T-test with Welch’s correction: % Open-Arm Time t_11.5_=2.841, p=0.0155; Open-Arm Entries t_12.6_=2.842, p=0.0143]. Female rats, on the other hand, exhibited no effect of alcohol exposure on either measure (**Figure 5I**) [unpaired two-tailed T-test: % Open-Arm Time t_19_=0.849, p=0.407; Open-Arm Entries t_19_=0.353, p=0.728]. That female rats displayed somewhat lower open arm time relative to males raises the possibility of a floor effect precluding further effects of alcohol exposure. However, since number of open-arm entries and immobility episodes (**Supplemental Figure S2**) similarly did not differ we do not believe this to be the case, although future studies would benefit from additional assays of anxiety-like behavior.

## DISCUSSION

The present study sought to determine the extent to which chronic alcohol exposure altered inter-laminar glutamatergic tone onto deep-layer PCs in prelimbic PFC and whether such changes occurred equivalently between sexes. Among our observations, we report that chronic exposure results in significantly elevated glutamate release probability from Layer II-III pyramidal cells on deep-layer PCs in male, but not female, rats. Additional investigation revealed that both male and female rats display readily distinguishable deep-layer PC subpopulations (Type-A and –B) whose functional and behavioral significance are only beginning to be fully understood. Discriminating according to these subtypes, we further show that alcohol effects on top-down glutamatergic signaling appear specific to sub-cortically projecting Type A cells in male, but not female, rats. These changes co-occurred with elevated anxiety-like behavior with no apparent effects on working memory in male rats, implicating specific PFC PC subtype afferents in the expression of alcohol-induced anxiety-like behavior in males. Furthermore, the data suggests these cells are particularly vulnerable to the effects of alcohol. Interestingly, in addition to an apparent homeostatic recovery in top-down glutamate signaling onto deep-layer PCs, female rats also exhibited enhanced performance on the NMS T-maze working memory task that further underscores the necessity of including sex as a relevant biological variable in the study of alcohol use disorders.

### Diversity in Deep-Layer Cortical Pyramidal Cells

The stratified layers of cortex host a rich milieu of excitatory and inhibitory neurons that adopt highly specialized functional roles within a cortical column [37-39], permitting significant computational power. We and others have investigated the effects of chronic ethanol exposure on a number of these cell subtypes, finding consistent adaptations that bias frontal cortical regions toward a hyperexcitable state [12, 13, 40]. While a number of studies have investigated excitatory signaling within cortex in a variety of disease models, most neglect to parse pyramidal cell subtype-specific changes despite a wealth of anatomic, genetic, and physiological data demonstrating at least two apparent subtypes [32, 41, 42].

Type-A and -B PC subtypes are anatomically distinguishable by projection destination, with Type-As preferentially distributing afferents to mid- and hind-brain structures via the pyramidal tracts while Type-B cells predominantly project intra-telencephalically [31, 33, 42]. Functionally, Type-A neurons also display a characteristic hyperpolarization-activated cyclic nucleotide gated channel (HCN)-mediated voltage sag when hyperpolarizing current is applied that is notably modest in Type-B cells, and exhibit a distinct facilitating response to excitatory signaling from superficial-layer pyramidal cells (**Figure 2A-B**) [30, 32]. Further characterization of this phenomenon demonstrated an apparent predisposition for Type-A cells as coincidence detectors of incoming excitation due to reliable facilitating synaptic responses over a narrow temporal window (**Figure 2D-E**) [41]. Reducing the response selectivity of these neurons, for instance via a generalized enhancement of presynaptic glutamate release probability (**Figure 3A**), would diminish the response specificity of these cells and reduce their capacity to filter out otherwise “noisy” information. Given the preferential interconnectivity of these cells with regions like the peri-aqueductal gray (PAG), shown to regulate fear and anxiety-like behaviors [42, 43], it is reasonable to expect that doing so would elicit elevations in such behaviors to otherwise innocuous stimuli (e.g. **Figure 5H**). In contrast, Type-B cells integrate excitatory inputs from local PCs as well as regions including hippocampus over a wide temporal window that includes the gamma frequency range [41]. This latter phenomenon in particular suggests a specific role for these cells in the generation and maintenance of reverberant gamma oscillations characteristic of working memory [44, 45], and indeed clinical evidence demonstrates significant correlations between aberrant gamma band patterns and impairments in working memory [46]. Based on these phenomena, it is tempting to postulate that alcohol-induced changes in Type-A orm -B cell activity would elicit concomitant changes in anxiety-like behavior or working memory performance respectively, and the present observations off some insight into the validity of this schema.

### Preferential Alcohol Effects on PFC-Dependent Behaviors Between Sexes

Indeed, we report robust changes in Type-A cell function that specifically co-occur with elevated anxiety-like behavior in males, suggesting these cells influence expression of anxiety-like behavior and are preferentially affected by alcohol exposure. Consistent with this scenario are recent observations showing that PAG-projecting PFC neurons exhibit physiological properties characteristic of Type-A cells, and mediate the expression of both fear-conditioned and alcohol drinking behaviors, appearing to function as a behavioral “brake” on consumption [42]. Extended alcohol exposure, though, elicits a hyperactive state in Type-A cells during acute withdrawal that coincides with heightened anxiety-like behavior, and one interpretation may be that overactive Type-A cells projecting to regions like PAG are promoting expression of these behaviors. Indeed, such a scenario is consistent with observations that specifically inactivating the prelimbic PFC diminishes the expression of these behaviors [36]. That we observe no effect of ethanol on either Type-A cell activity or anxiety-like behavior in females supports this assertion, particularly in light of other evidence that females appear resilient to PFC-mediated elevations in anxiety-like behavior after ethanol exposure [11]. Alternatively, we have previously reported that PFC PCs projecting to CeA display reduced inhibitory tone, and while it is not presently known if these cells are Type-A or Type-B, their heightened downstream signaling to CeA is nevertheless consistent with enhance expression of anxiety-like behavior in males [13]. Indeed, others report that increased PFC signaling into the adjacent basolateral amygdala (BLA) during acute withdrawal is necessary and sufficient to elicit such behaviors [34], though again it is unclear from which PC subclass these terminals emanate. Considered together, it seems reasonable to hypothesize that hyperactivity specifically within Type-A cells diminishes the specificity of PFC top-down control over the expression of fear/anxiety-like behaviors, contributing to negative affect states characteristic of ethanol dependence. This raises the question as to what predisposes this cell population to the effects of chronic alcohol exposure, both pre- and post-synaptically. That deep-layer Type-A and –B PCs show apparent subtype-specific genetic markers suggests the possibility of selectively targeting one or the other population, and doing so represents fertile ground for additional study [47, 48].

Though the specific mechanism(s) governing alcohol effects on glutamate release probability onto PC cells remain unclear, based on observations modulating extracellular calcium (**Figure 3C-D**) it is likely such changes reflect alterations in calcium-dependent presynaptic processes including voltage gated calcium channel (VGCC) expression and/or function, which is further supported by data showing no effect of picrotoxin on this phenomenon (**Figure 1G-I**). Reports demonstrating that adaptations in VGCCs as well as other proteins integral to presynaptic vesicular release govern enhanced glutamate release probability after chronic ethanol exposure support this hypothesis [49, 50]. Indeed, it is also not presently known whether inputs to Type-A/-B cells differentially express VGCC subtypes, representing an additional avenue for investigation. Another open question is the degree to which concomitant changes in glutamate and GABA tone on deep-layer PCs [13, 26] specifically drive altered excitability in Type-A cells (**Figure 3G**). Similar elevations in PC excitability have been observed in lateral orbital frontal cortex after chronic alcohol exposure due to reductions in small conductance calcium-activated potassium (SK) channel-mediated conductance [40], and differential expression or activity of these channels may contribute to our observations in PFC, warranting further investigation. Though not addressed in the present work, chronic alcohol exposure has also been shown to potently alter monoamine signaling in cortex [51, 52], and as Type-A PCs appear especially sensitive to regulation by neuromodulators [30, 31], it would be of substantial interest to evaluate how such changes coordinate with alcohol effects on synaptic scaling to influence excitability between PC subtypes.

As a defining feature of Type-A cells, it is somewhat curious we did not observe any apparent effects of ethanol exposure on HCN channel activity in these cells, particularly given recent reports of significant modulation of these channels by alcohol exposure [19, 24]. Observations by Salling, Skelly [24], though, were conducted using an adolescent alcohol exposure model that, due to extensive developmental regulation of HCNs during adolescence [23], likely underlies incongruent findings between their report and this paper. As well, our measurements are likely only detecting somatic or proximal dendrite-expressed HCNs, the possibility these channels are undergoing extensive spatial remodeling or functional modification cannot be ruled out. In any case, it is notable that we again observe higher V_H_ in female cells than males, consistent with recent findings by our laboratory showing similarly elevated HCN-mediated activity in Martinotti interneurons and HCN1 protein expression in PFC [19]. Importantly, that study further showed differential sex-specific changes in both Martinotti cell excitability and HCN-mediated current that could explain female-specific alcohol-facilitated working memory performance despite no apparent effects on Type-B PCs. Such a schema is consistent with previous observations showing somatostatin-positive interneurons, which include Martinotti cells, are intimately involved in mediating working memory [53]. That elevated basal levels of HCNs in PFC co-occur with somewhat higher performance during NMS T-maze training in females compared to males (**Figure 5B,E**) additionally conforms to human data showing similarly higher performance on working memory tasks in women, further supporting the conclusion that cortical circuits and behaviors cannot be assumed equivalent between sexes [54, 55]. Though sex-specific alterations in hippocampal input to PFC after alcohol exposure may also underlie disparate effects on working memory [35], given the prominent role of HCNs in cortical circuit function and working memory [23, 41, 45], investigating differential innate and ethanol-induced expression patterns of cortical HCNs on working memory between sexes is a particularly compelling avenue for future investigation.

## Conclusions

In sum, the present work demonstrates evidence that chronic alcohol exposure promotes differential adaptations in both prefrontal cortical cell excitability and PFC-dependent behaviors between male and female rats that further underscores the need to consider sex as a relevant biological variable in studies of AUDs. Additionally, we show that distinct deep-layer PC subpopulations are apparent in both males and females, and display complex adaptations that warrant further investigation. Future studies will seek to discern the role of these cells in mediating alcohol-induced perturbations in select cortical microcircuits to elicit specific aberrant behaviors.

## AUTHOR CONTRIBUTIONS

Study design, data collection and analysis, and manuscript preparation were conducted by BAH. TKO and GB assisted with EPM experiments shown in **Figure 5**. MAH and ALM assisted with interpretation of results and preparation of the manuscript.

## REFERENCES

1. World Health Organization, Global Status Report on Alcohol and Health 2014. Web, 2014.

2. Rehm, J., The risks associated with alcohol use and alcoholism. Alcohol Res Health, 2011. 34(2): p. 135–43.

3. Anthony, J.C., L.A. Warner, and R.C. Kessler, Comparative epidemiology of dependence on tobacco, alcohol, controlled substances, and inhalants: Basic findings from the National Comorbidity Survey. Experimental and Clinical Psychopharmacology, 1994. 2(3): p. 244–268.

4. Rehm, J., et al., Burden of disease associated with alcohol use disorders in the United States. Alcohol Clin Exp Res, 2014. 38(4): p. 1068–77.

5. Kushner, M.G., K. Abrams, and C. Borchardt, The relationship between anxiety disorders and alcohol use disorders: a review of major perspectives and findings. Clin Psychol Rev, 2000. 20(2): p. 149–71.

6. Koob, G.F. and N.D. Volkow, Neurocircuitry of addiction. Neuropsychopharmacology, 2010. 35(1): p. 217–38.

7. Grant, B.F., et al., Epidemiology of DSM-5 Alcohol Use Disorder: Results From the National Epidemiologic Survey on Alcohol and Related Conditions III. JAMA Psychiatry, 2015. 72(8): p. 757– 66.

8. Clayton, J.A. and F.S. Collins, Policy: NIH to balance sex in cell and animal studies. Nature, 2014. 509(7500): p. 282–3.

9. Nolen-Hoeksema, S. and L. Hilt, Possible contributors to the gender differences in alcohol use and problems. J Gen Psychol, 2006. 133(4): p. 357–74.

10. Morales, M., M.M. McGinnis, and B.A. McCool, Chronic ethanol exposure increases voluntary home cage intake in adult male, but not female, Long-Evans rats. Pharmacol Biochem Behav, 2015. 139(Pt A): p. 67–76.

11. Morales, M., et al., Chronic Intermittent Ethanol Exposure Modulation of Glutamatergic Neurotransmission in Rat Lateral/Basolateral Amygdala is Duration-, Input-, and Sex-Dependent. Neuroscience, 2018. 371: p. 277–287.

12. Bohnsack, J.P., et al., Histone deacetylases mediate GABAA receptor expression, physiology, and behavioral maladaptations in rat models of alcohol dependence. Neuropsychopharmacology, 2018. 43(7): p. 1518–1529.

13. Hughes, B.A., et al., Chronic Ethanol Exposure and Withdrawal Impairs Synaptic GABAA Receptor-Mediated Neurotransmission in Deep Layer Prefrontal Cortex. Alcohol Clin Exp Res, 2019. 43(5): p. 822–832.

14. Devaud, L.L., D.B. Matthews, and A.L. Morrow, Gender impacts behavioral and neurochemical adaptations in ethanol-dependent rats. Pharmacology, Biochemistry and Behavior, 1999. 64(4): p. 841–849.

15. Follesa, P. and M.K. Ticku, Chronic ethanol-mediated up-regulation of the N-methyl-D-aspartate receptor polypeptide subunits in mouse cortical neurons in culture. J Biol Chem, 1996. 271(23): p. 13297–9.

16. Carpenter-Hyland, E.P., J.J. Woodward, and L.J. Chandler, Chronic ethanol induces synaptic but not extrasynaptic targeting of NMDA receptors. J Neurosci, 2004. 24(36): p. 7859–68.

17. Devaud, L.L. and A.L. Morrow, Gender-selective effects of ethanol dependence on NMDA receptor subunit expression in cerebral cortex, hippocampus and hypothalamus. European Journal of Pharmacology, 1999. In Press.

18. Nimitvilai, S., M.F. Lopez, and J.J. Woodward, Sex-dependent differences in ethanol inhibition of mouse lateral orbitofrontal cortex neurons. Addict Biol, 2020. 25(1): p. e12698.

19. Hughes, B.A., et al., Chronic ethanol exposure alters prelimbic prefrontal cortical Fast-Spiking and Martinotti interneuron function with differential sex specificity in rat brain. Neuropharmacology, 2020. 162: p. 107805.

20. Joffe, M.E., D.G. Winder, and P.J. Conn, Contrasting sex-dependent adaptations to synaptic physiology and membrane properties of prefrontal cortex interneuron subtypes in a mouse model of binge drinking. Neuropharmacology, 2020. 178: p. 108126.

21. Besheer, J., et al., The effects of repeated corticosterone exposure on the interoceptive effects of alcohol in rats. Psychopharmacology (Berl), 2012. 220(4): p. 809–22.

22. Kotlinska, J. and M. Bochenski, The influence of various glutamate receptors antagonists on anxiety-like effect of ethanol withdrawal in a plus-maze test in rats. Eur J Pharmacol, 2008. 598(1-3): p. 57–63.

23. Wang, M., et al., Alpha2A-adrenoceptors strengthen working memory networks by inhibiting cAMP-HCN channel signaling in prefrontal cortex. Cell, 2007. 129(2): p. 397–410.

24. Salling, M.C., et al., Alcohol Consumption during Adolescence in a Mouse Model of Binge Drinking Alters the Intrinsic Excitability and Function of the Prefrontal Cortex through a Reduction in the Hyperpolarization-Activated Cation Current. J Neurosci, 2018. 38(27): p. 6207– 6222.

25. DeHart, W.B. and B.A. Kaplan, Applying mixed-effects modeling to single-subject designs: An introduction. J Exp Anal Behav, 2019.

26. Pleil, K.E., et al., Effects of chronic ethanol exposure on neuronal function in the prefrontal cortex and extended amygdala. Neuropharmacology, 2015. 99: p. 735–49.

27. Fioravante, D., et al., Calcium-dependent isoforms of protein kinase C mediate posttetanic potentiation at the calyx of Held. Neuron, 2011. 70(5): p. 1005–19.

28. Thanawala, M.S. and W.G. Regehr, Presynaptic calcium influx controls neurotransmitter release in part by regulating the effective size of the readily releasable pool. J Neurosci, 2013. 33(11): p. 4625–33.

29. Gioia, D.A., N.J. Alexander, and B.A. McCool, Differential Expression of Munc13-2 Produces Unique Synaptic Phenotypes in the Basolateral Amygdala of C57BL/6J and DBA/2J Mice. J Neurosci, 2016. 36(43): p. 10964–10977.

30. Dembrow, N.C., R.A. Chitwood, and D. Johnston, Projection-specific neuromodulation of medial prefrontal cortex neurons. J Neurosci, 2010. 30(50): p. 16922–37.

31. Gee, S., et al., Synaptic activity unmasks dopamine D2 receptor modulation of a specific class of layer V pyramidal neurons in prefrontal cortex. J Neurosci, 2012. 32(14): p. 4959–71.

32. Lee, A.T., et al., Pyramidal neurons in prefrontal cortex receive subtype-specific forms of excitation and inhibition. Neuron, 2014. 81(1): p. 61–8.

33. Reiner, A., et al., Differential morphology of pyramidal tract-type and intratelencephalically projecting-type corticostriatal neurons and their intrastriatal terminals in rats. J Comp Neurol, 2003. 457(4): p. 420–40.

34. McGinnis, M.M., et al., Chronic Ethanol Differentially Modulates Glutamate Release from Dorsal and Ventral Prefrontal Cortical Inputs onto Rat Basolateral Amygdala Principal Neurons. eNeuro, 2020. 7(2).

35. Bygrave, A.M., et al., Hippocampal-prefrontal coherence mediates working memory and selective attention at distinct frequency bands and provides a causal link between schizophrenia and its risk gene GRIA1. Transl Psychiatry, 2019. 9(1): p. 142.

36. Sierra-Mercado, D., N. Padilla-Coreano, and G.J. Quirk, Dissociable roles of prelimbic and infralimbic cortices, ventral hippocampus, and basolateral amygdala in the expression and extinction of conditioned fear. Neuropsychopharmacology, 2011. 36(2): p. 529–38.

37. Silberberg, G. and H. Markram, Disynaptic inhibition between neocortical pyramidal cells mediated by Martinotti cells. Neuron, 2007. 53(5): p. 735–46.

38. Owen, S.F., J.D. Berke, and A.C. Kreitzer, Fast-Spiking Interneurons Supply Feedforward Control of Bursting, Calcium, and Plasticity for Efficient Learning. Cell, 2018. 172(4): p. 683–695 e15.

39. Naka, A. and H. Adesnik, Inhibitory Circuits in Cortical Layer 5. Front Neural Circuits, 2016. 10: p. 35.

40. Nimitvilai, S., et al., Chronic Intermittent Ethanol Exposure Enhances the Excitability and Synaptic Plasticity of Lateral Orbitofrontal Cortex Neurons and Induces a Tolerance to the Acute Inhibitory Actions of Ethanol. Neuropsychopharmacology, 2016. 41(4): p. 1112–27.

41. Dembrow, N.C., B.V. Zemelman, and D. Johnston, Temporal dynamics of L5 dendrites in medial prefrontal cortex regulate integration versus coincidence detection of afferent inputs. J Neurosci, 2015. 35(11): p. 4501–14.

42. Siciliano, C.A., et al., A cortical-brainstem circuit predicts and governs compulsive alcohol drinking. Science, 2019. 366(6468): p. 1008–1012.

43. Silva, C. and N. McNaughton, Are periaqueductal gray and dorsal raphe the foundation of appetitive and aversive control? A comprehensive review. Prog Neurobiol, 2019. 177: p. 33–72.

44. Roux, F., et al., Gamma-band activity in human prefrontal cortex codes for the number of relevant items maintained in working memory. J Neurosci, 2012. 32(36): p. 12411–20.

45. Constantinidis, C., et al., Persistent Spiking Activity Underlies Working Memory. J Neurosci, 2018. 38(32): p. 7020–7028.

46. Senkowski, D. and J. Gallinat, Dysfunctional prefrontal gamma-band oscillations reflect working memory and other cognitive deficits in schizophrenia. Biol Psychiatry, 2015. 77(12): p. 1010–9.

47. Molyneaux, B.J., et al., Novel subtype-specific genes identify distinct subpopulations of callosal projection neurons. J Neurosci, 2009. 29(39): p. 12343–54.

48. Molnar, Z. and A.F. Cheung, Towards the classification of subpopulations of layer V pyramidal projection neurons. Neurosci Res, 2006. 55(2): p. 105–15.

49. Gioia, D.A., N. Alexander, and B.A. McCool, Ethanol Mediated Inhibition of Synaptic Vesicle Recycling at Amygdala Glutamate Synapses Is Dependent upon Munc13-2. Front Neurosci, 2017. 11: p. 424.

50. Varodayan, F.P., et al., Alcohol Dependence Disrupts Amygdalar L-Type Voltage-Gated Calcium Channel Mechanisms. J Neurosci, 2017. 37(17): p. 4593–4603.

51. Nimitvilai, S., et al., Ethanol Dependence Abolishes Monoamine and GIRK (Kir3) Channel Inhibition of Orbitofrontal Cortex Excitability. Neuropsychopharmacology, 2017. 42(9): p. 1800– 1812.

52. Trantham-Davidson, H., et al., Binge-Like Alcohol Exposure During Adolescence Disrupts Dopaminergic Neurotransmission in the Adult Prelimbic Cortex. Neuropsychopharmacology, 2017. 42(5): p. 1024–1036.

53. Kim, D., et al., Distinct Roles of Parvalbumin- and Somatostatin-Expressing Interneurons in Working Memory. Neuron, 2016. 92(4): p. 902–915.

54. Rahman, Q., S. Abrahams, and F. Jussab, Sex differences in a human analogue of the Radial Arm Maze: the “17-Box Maze Test”. Brain Cogn, 2005. 58(3): p. 312–7.

55. Reed, J.L., et al., Sex differences in verbal working memory performance emerge at very high loads of common neuroimaging tasks. Brain Cogn, 2017. 113: p. 56–64.

