## Supplementary Material for "Sex- and Subtype-Specific Adaptations in Excitatory Signaling Onto Deep-Layer Prelimbic Cortical Pyramidal Neurons After Chronic Alcohol Exposure"

**SUPPLEMENTAL METHODS**

*Electrophysiological Recordings*

Repeated pulse responses (RPRs) shown in **Figures 1-4** are normalized responses to the first pulse of each individual recording (e.g. baseline or picrotoxin/low-Ca^2+^). Addition of picrotoxin (100 µM) resulted in epileptiform activity with >10 Hz stimulation and was therefore only used for experiments shown in **Figure 1**. Nevertheless, application of picrotoxin did not affect amplitudes or half-widths (data not shown), demonstrating evoked currents were predominantly AMPA receptor-mediated. In experiments shown in **Figures 2-4**, after obtaining a baseline recording, the amplifier was switched to current-clamp mode and membrane voltage responses to a series of hyper- and depolarizing current steps obtained. On returning to voltage-clamp, cells were then perfused with drug (e.g. picrotoxin, low-Ca^2+^ aCSF, etc) for ~5 minutes followed by another series of stimulation frequency-response recordings. Access resistance was monitored throughout the experiment, and any recordings exceeding 25 MΩ were terminated. For **Figures 2B-C**, data were calculated by adding together the peak “sag” in response to -250 pA current injection, the peak “overshoot” when current was removed, and the peak sag following removal of +250 pA current injection. The sum was then normalized to the steady-state average V_R_ of the -250 pA current injection (e.g. last 50 ms) and expressed as a percent. For all electrophysiological experiments, individual cells were defined as the unit of determination, recorded from a minimum of 3 rats per group.

*Elevated Plus Maze (EPM)*

Rats were brought into a partially-lit (~20 lux) procedure room approximately 20 minutes prior to testing to acclimate, then placed into the EPM apparatus (Stoelting Co.; Wood Dale, IL) and permitted to freely explore for 5 minutes. Behavior was recorded and analyzed off-line using ANY-Maze tracking software (Stoelting Co., v4.82) to measure % of time spend in open arms, open arm entries, immobile episodes (defined as periods of immobility longer than 10 s), and total distance traveled.

*Delayed Non-Match-to-Sample (NMS) T-Maze*

Briefly, all animals were food restricted to ~95% of free feeding weight for 2 days prior and during NMS T-Maze training. Training on the task consisted of an initial forced-choice phase followed by an open choice phase 30 seconds after. Specifically, during the forced trial one arm is blocked (e.g. the left) and in the other (e.g. right) a sucrose reward (chocolate) is placed. Following entry (defined as all 4 legs within the plane of the arm) and reward consumption the rat is placed in a holding chamber, and the door to the other arm is removed and a sucrose reward placed at the end. After the latency period (30 s), the rat must then choose either the new correct (rewarded) arm or the incorrect (unrewarded) arm and is blocked in for 10 seconds. Rats performed this sequence 10 times per day with the forced arm alternated randomly such that five trials begin left or right. At the end of training the animals were removed from food restriction and 24 hours after underwent water or ethanol exposure (15 days once-daily 5.0 g/kg ethanol, i.g.). 2 days prior to final exposure rats were again food restricted, and starting 24 hours after the last administration underwent 3 days of NMS T-maze testing. During this phase, the animal performed the task 6 times with increasing latencies (10, 60, and 120 seconds; 2 times each in random order) between the forced and choice trials, with the forced arm counterbalanced to include left and right for each latency trial. 24 hours after the final test day rats were sacrificed and brains collected.

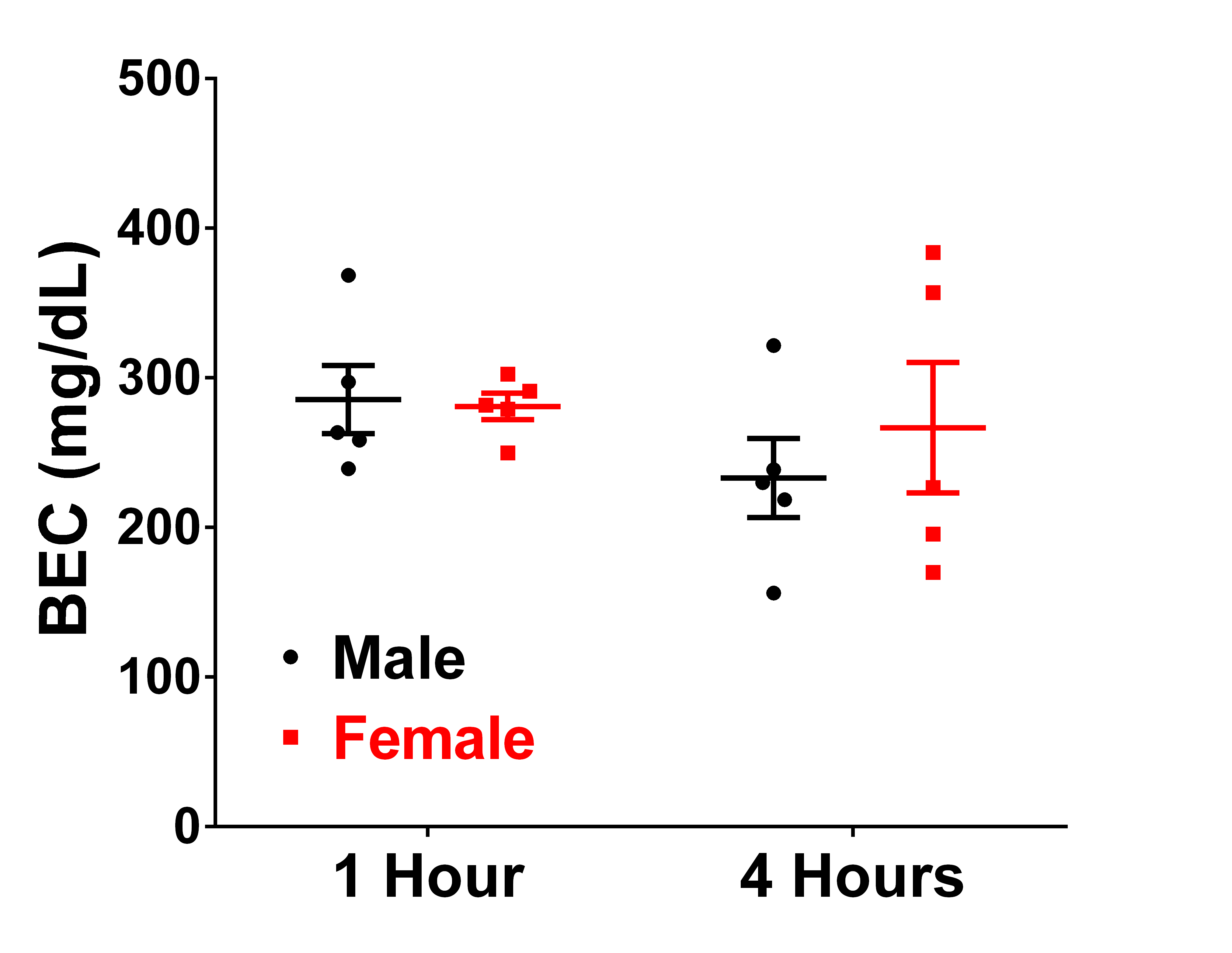

**Supplemental Figure S1:** 5.0 g/kg (25% v/v) ethanol gavage in naïve male and female rats elicits peak BECs of ~300 mg/dL after 1 hour. BECs were analyzed in a separate group of naïve rats 4 hours post-gavage, finding similar BECs indicative of sustained intoxication.

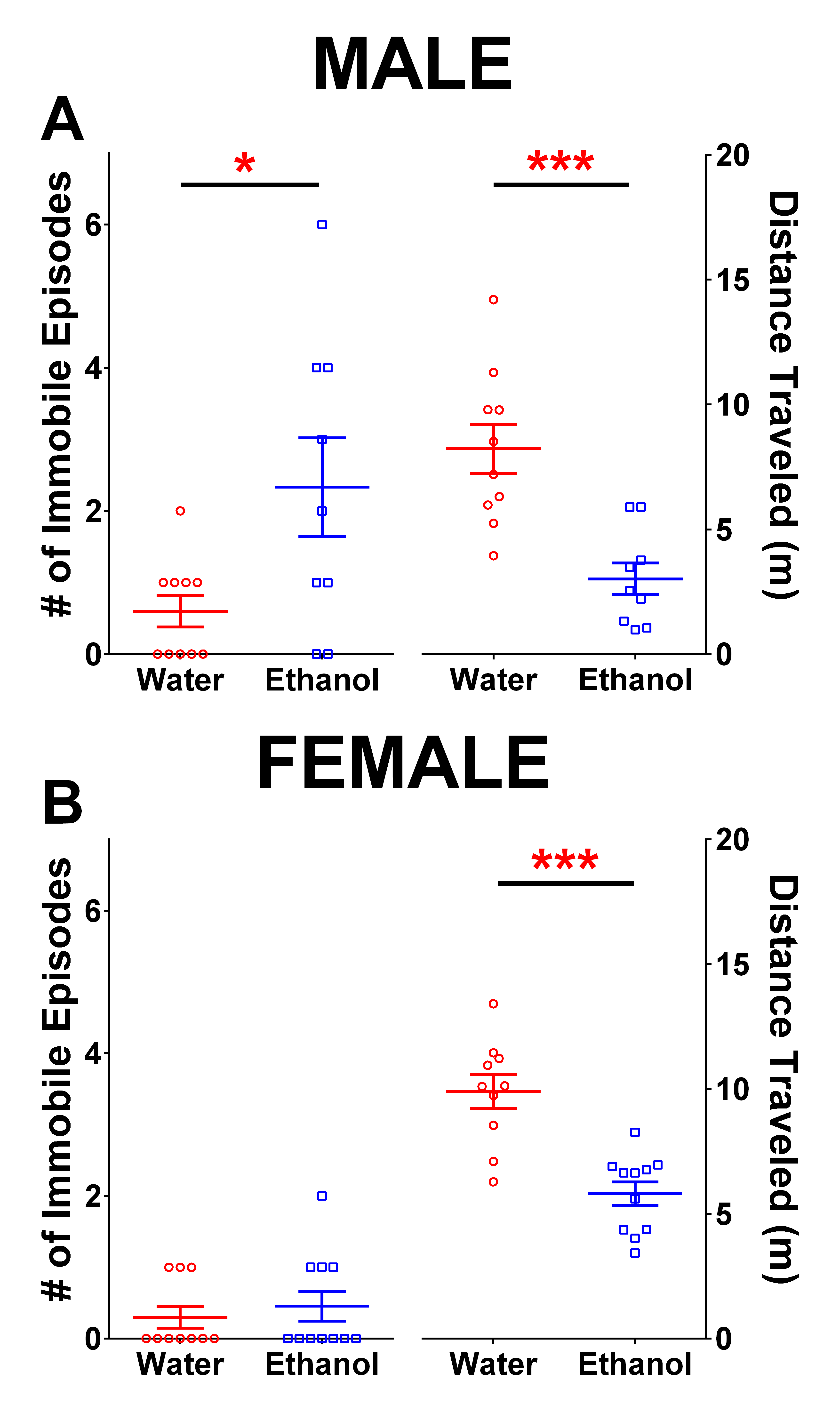

**Supplemental Figure S2:** Number of immobility episodes and total distance traveled measures from EPM behavioral assay. A) Male ethanol exposed rats displayed a significant increase in the number of immobile periods [unpaired two-tailed T-test with Welch’s correction: t_9.65_=2.401, p=0.038] and corresponding decrease in total distance traveled [t_15.12_=4.451, p=0.0005]. B) Ethanol exposure did not alter number of immobile episodes [unpaired two-tailed T-test: t_19_=0.59, p=0.562] but did significantly reduce total distance traveled [t_19_=5.07, p<0.0001].

| **Cell** | **C_MEM_ (pF)** | **R_MEM_ (MΩ)** | **RMP (mV)** | **V_H_ (mV)** | **Cell** | **C_MEM_ (pF)** | **R_MEM_ (MΩ)** | **RMP (mV)** | **V_H_ (mV)** |
| --- | --- | --- | --- | --- | --- | --- | --- | --- | --- |
| **Male** | **104.1 ± 4.1 (W)**  **96.7 ± 3.9 (E)** | **70.6 ± 2.1 (W)**  **76.3 ± 3.3 (E)** | **-66.7 ± 1.0 (W)**  **-67.6 ± 0.6 (E)** | **11.7 ± 0.5 (W)**  **13.1 ± 0.6 (E)** | **Male** | **86.6 ± 3.1 (W)**  **89.3 ± 4.3 (E)** | **91.1 ± 4.3 (W)**  **81.8 ± 5.8 (E)** | **-73.3 ± 0.9 (W)**  **-69.7 ± 1.9 (E)** | **8.3 ± 0.6 (W)**  **7.7 ± 0.5 (E)** |
| **Type A** |  |  |  |  | **Type B** |  |  |  |  |
| **Female** | **97.9 ± 7.2 (W)**  **112.8 ± 6.9 (E)** | **89.7 ± 7.7 (W)**  **73.8 ± 4.0 (E)** | **-67.1 ± 0.8 (W)**  **-66.4 ± 1.3 (E)** | **15.0 ± 0.6 (W)**  **17.4 ± 1.2 (E)** | **Female** | **90.2 ± 5.4 (W)**  **89.7 ± 3.2 (E)** | **84.1 ± 4.0 (W)**  **93.2 ± 5.6 (E)** | **-70.3 ± 1.4 (W)**  **-71.4 ± 1.3 (E)** | **10.3 ± 0.9 (W)**  **10.7 ± 1.3 (E)** |
| **Type A** |  |  |  |  | **Type B** |  |  |  |  |

**Table 1: Electrophysiological properties of Type-A and –B deep-layer Pyramidal Neurons in prelimbic PFC**

Data presented as mean±SEM. No significant differences were observed between water- or ethanol-exposed groups among the four cell populations.
